# Large-scale mutational analysis uncovers molecular mechanisms governing dual RNA functions in transposons

**DOI:** 10.64898/2026.03.11.711047

**Authors:** Edan E. Mortman, Samuel H. Sternberg

## Abstract

Transposons are among the most abundant mobile genetic elements in nature. IStrons are a unique class of transposons, encoding a transposase for transposon mobility, an RNA-guided nuclease for maintenance, and a self-splicing group I intron for element removal from host mRNA. However, it is unclear how a single polynucleotide sequence balances these distinct biochemical functions. Here we employed pooled library mutagenesis coupled with high-throughput sequencing to systematically dissect the molecular determinants of IStron transposition, RNA-guided DNA cleavage, and self-splicing. We found that the terminal trinucleotide of the transposon right end is constrained by all three functions, identifying a molecular convergence point. Cross-assay comparisons revealed that most variants maintained or lost activity across multiple assays simultaneously. However, a subset selectively retained one activity while losing another, revealing an antagonism between DNA cleavage and splicing governed by guide RNA structural stability. Increased GC content at the base of the guide RNA 3’ terminal stem-loop abolished splicing while maintaining DNA cleavage, and the stably folded guide RNA sterically occludes alternative splice sites, ensuring splicing accuracy across variable flanking contexts. Thus, IStron transcripts face an inherent trade-off between guide RNA maturation and splicing, with RNA structural stability as the primary determinant of pathway choice.

## INTRODUCTION

Transposable elements use RNAs for diverse functions in their life cycle, from replicative intermediates in retrotransposon (1) to guide RNAs that direct site-specific transposition by CRISPR-associated transposons (CASTs) (2, 3) and IS110 (4) elements. Insertion Sequence 605 (IS605) and Insertion Sequence 607 (IS607) families similarly employ critical, transposon-encoded non-coding RNAs, though their primary functions are to guide endonuclease-mediated target DNA cleavage for element retention (5–10). Although IS605 and IS607 elements encode distinct transposases for mobilization, termed TnpA, they both encode the same accessory protein, TnpB. Recent work has shown that TnpB is an endonuclease that forms a ribonucleoprotein complex with an OMEGA guide RNA (ωRNA) encoded within the element boundaries. Intriguingly, a subset of these elements additionally encode a group I intron that spans the entire element from the left end (LE) to the right end (RE), and are thus termed “IStrons” (11–14). These self-splicing ribozymes have the capacity to remove themselves from interrupted transcripts, potentially mitigating the fitness cost of transposon insertion.

In IS607 family IStrons, TnpA-mediated excision removes the element from the host genome, generating circular DNA intermediates that can transpose to new genomic locations (10). Critically, this excision produces scarless donor joints that regenerate the original genomic sequence, placing elements at risk of copy number loss when excision frequency outpaces integration at new target sites. The TnpB-ωRNA complex counteracts this loss by specifically recognizing and cleaving the scarless donor joint through RNA-guided DNA cleavage. This recognition requires a transposon-adjacent motif (TAM) flanking the guide-complementary target sequence, enabling TnpB to discriminate between sites containing the element and those lacking it. The resulting double-strand break promotes RecA-dependent homologous recombination, leading to IStron retention at the original site (10). Thus, the TnpB-ωRNA complex plays a crucial role in promoting the spread of IStrons by counteracting transposon loss.

The IStron RNA can also fold into a specific group I intron structure, which facilitates autonomous self-splicing through a two-step transesterification mechanism that requires only Mg^2+^ and an exogenous guanosine triphosphate (GTP) (15–17). By splicing out of interrupted transcripts and rejoining flanking exonic sequences, group I introns restore interrupted gene sequence and function (18)-(19). For IStrons, this ability has the potential to mitigate the fitness costs typically associated with insertional mutagenesis. However, folding into a group I intron creates an inherent conflict: the intron 3’ splice site occurs precisely within the ωRNA scaffold, such that splicing severs the scaffold from the guide sequence, thus inactivating the ωRNA (10) (**Fig. 1A**). This conflict is compounded by the fact that the IStron right end (RE) contains overlapping sequence determinants that must simultaneously satisfy three distinct biochemical requirements: recognition by TnpA at the DNA level for excision, folding into a functional ωRNA scaffold for TnpB-mediated cleavage, and proper positioning of the 3’ splice site for group I intron catalysis. Self-splicing therefore offers the potential advantage of mitigating insertional mutagenesis by restoring interrupted gene function, while reducing the pool of functional ωRNA molecules available for TnpB-mediated retention. This raises the question of how IStrons balance these competing molecular functions, encoded within a shared sequence, to maintain element survival and host viability.

**Figure 1.**
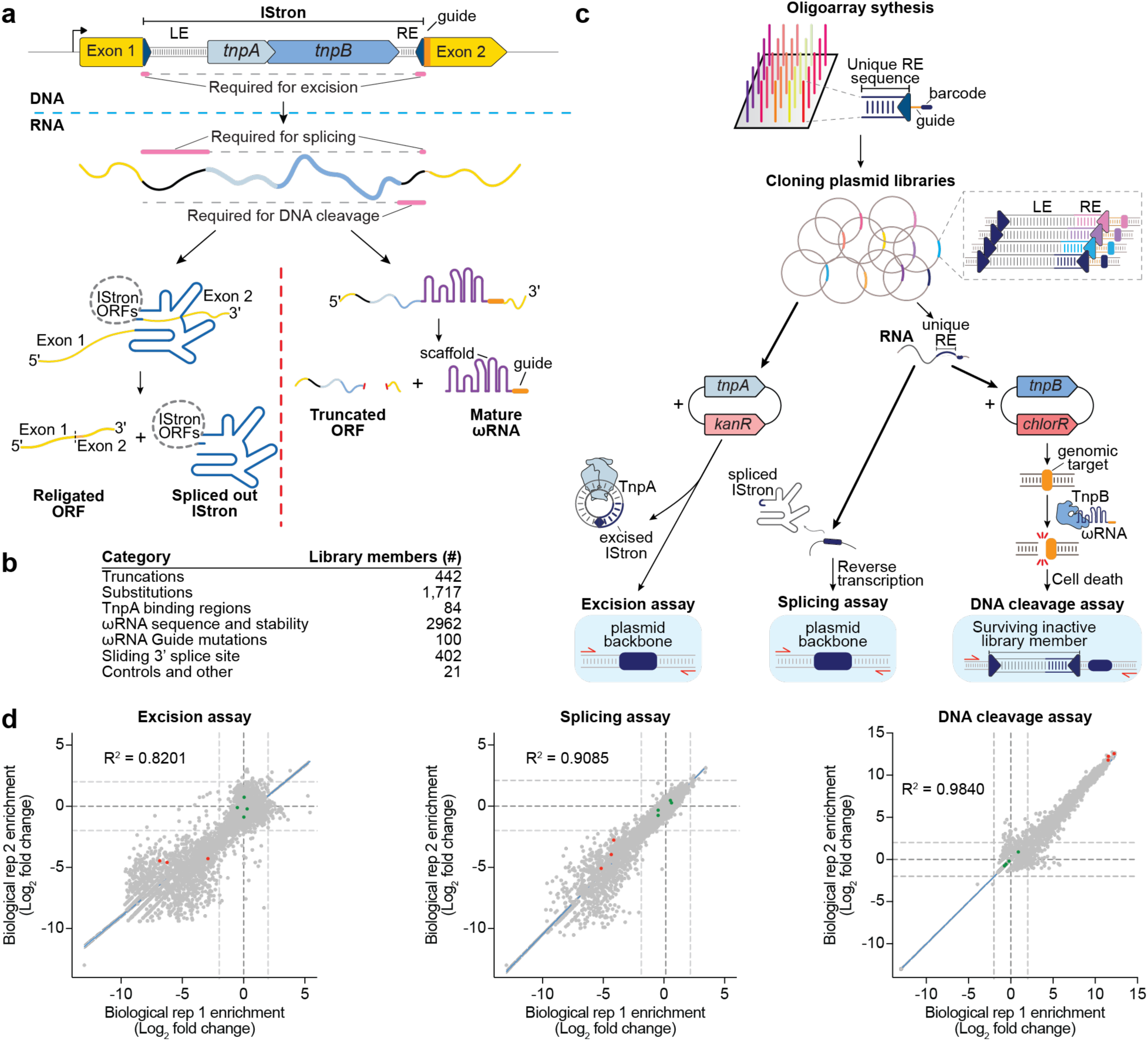
Systematic mutagenesis library for dissecting IStron functional determinants. **a)** Schematic of IStron organization showing overlapping sequence requirements for TnpA-mediated excision, TnpB-ωRNA DNA cleavage, and group I intron splicing (nt portions depicted in pink). DNA level (top) depicts IStron boundaries, containing the IStron left- and right-ends (LE and RE) and TnpA and TnpB coding regions, inserted into a host gene, creating two exons. RNA level (bottom) shows competing pathways: group I intron splicing that religates bacterial ORF and splices out the IStron, versus TnpB-ωRNA complex formation for DNA cleavage. **b)** Categories and abundances of library variants. **c)** Library construction workflow. Oligoarray synthesis generated pooled variants with unique barcodes, followed by cloning into a pooled plasmid library. Parallel assays were performed: excision with TnpA co-expression, splicing via RNA extraction and reverse transcription, and DNA cleavage with TnpB co-expression leading to cell death for functional ωRNAs. **d)** Biological replicate correlation for each library assay. Scatter plots show log_2_-fold change of library members normalized to wild-type for excision, splicing, and DNA cleavage assays. WT IStron library members are in green; representatives of each scrambled RE negative control are in red. Light gray dotted lines denote the log_2_-fold change of 2 and -2; dark gray dotted lines denote the log_2_ fold change of 0. R^2^ values from linear regression are shown for each assay.

Here, we systematically dissected the sequence determinants governing TnpA-mediated excision, TnpB–ωRNA mediated DNA cleavage, and group I intron-mediated self-splicing, to determine how IStrons achieve this intricate balance. By designing a pooled oligonucleotide library encoding thousands of IStron variants, we performed parallel evaluation of each variant’s performance across all three molecular functions. This approach revealed that IStrons have evolved to maintain molecular constraints, such as a terminal trinucleotide sequence, that are required across all three IStron functions. By comparing library variants across assays, we showed that splicing is subordinate to ωRNA formation and that splicing is easily lost and occurs less readily than DNA cleavage. Finally, by focusing on splicing sequence determinants, we found that the 3’ exon plays a major role in splicing efficiency and splice site selection, demonstrating that external sequence features can also affect IStron life cycle outcomes. Our results demonstrate how transposon-encoded RNAs have evolved to balance mutually exclusive activities, prioritizing element survival while also accommodating host fitness mitigation.

## MATERIALS AND METHODS

### Design, cloning and analysis of pooled IStron plasmid library

Library members were designed using a combination of custom scripts for systematic variants such as the truncation and substitution datasets, random sequence generation for degenerate positions, and manual curation for specific constructs of interest.

The RE variants within the IStron plasmid library were first generated as 300-nt single-stranded pooled oligos (Twist Bioscience). All variants are listed in Supplementary Table 3. Variant design was either automated using custom scripts or designed by hand in spreadsheets. 10 ng of oligoarray library DNA was amplified by PCR for 12 cycles in 50 µl reactions using Kapa DNA polymerase (Roche) and primers that bind to all library members, adding restriction enzyme digestion sites. All plasmids used in this study are listed in Supplementary Table S1, and oligos are listed in Supplementary Table S2. Amplicons were cleaned up using a MinElute Gel Extraction Kit (Qiagen) and eluted in 15 µl Mili-Q H_2_O. As the backbone vector, we used a plasmid encoding the WT mini-IStron sequence on a pCDF backbone. The backbone vector and library insert amplicons were digested using NcoI and NotI at 37^°^C for 1 h, gel purified, and ligated in 20 µl reactions with T4 DNA ligase (NEB) at 25^°^C for 30 min. Ligation reactions were cleaned up and eluted in 10 µl Mili-Q H_2_O (MinElute PCR Purification Kit), and then used to transform electrocompetent NEB 10-beta cells in four individual electroporation reactions according to the manufacturer’s protocol. After recovery (37^°^C for 2 h), transformed cells were plated on large 245 mm x 245 mm bioassay plates containing LB-agar with spectinomycin (100 µg ml^-1^) and grown for 24 h at 37^°^C. Plates were scraped to collect cells at a 100x coverage, and plasmid DNA was isolated using the Qiagen Plasmid Midi Kit.

10 ng of the isolated IStron plasmid DNA were used as input for PCR1 amplification with Kapa DNA polymerase (Roche) for 12 cycles, using specific primers that flank the variable RE position and barcode of each plasmid. Samples were cleaned up and eluted in 30 µl EB. 2 µl of the PCR1 product were amplified in 12 cycles for PCR2. Libraries were sequenced on an Element AVITI in paired-end mode with 150 cycles per end.

NGS data analysis was performed using custom Python scripts. Demultiplexed reads were filtered to remove reads that did not contain a perfect match to the 19 nt primer binding sequence at the 3’-terminus of the IStron end. Then, the 10 nt sequence directly downstream of the primer binding sequence was extracted, which encodes the barcode that uniquely identifies each library member. The number of reads containing each library member barcode was counted. If a read did not contain a barcode that matched a library member barcode, it was discarded. The relative abundance of each library member was then determined by dividing the barcode count of each library member by the total number of barcode counts. Coupling rates for the library input controls, one WT sequence with four different barcodes and three scrambled controls with four different barcodes each, were calculated as for **Figure S1F** (coupled reads divided by total reads per barcode). Across both biological replicates, the four wild-type barcodes coupled at 55.8 ± 3.2%. The twelve negative-control barcodes coupled at 39.5 ± 13.2% overall, with one control sequence coupling comparably to WT (∼53%) and the other two lower (∼31% and ∼34%). Because library-member abundance is quantified from barcode counts, and uncoupled reads are distributed stochastically across the library (**Fig. S1G**), this coupling difference adds noise but does not systematically bias the WT-normalized fold-change values (**Fig. S2A**).

### DNA cleavage assays with TnpB

In individual IStron variant experiments, *E. coli* strain BL21(DE3) was transformed with a pEffector plasmid containing a codon optimized TnpB (*Cbo*TnpB or *Cse*TnpB) on a pACYC backbone. Single colony isolates were selected to prepare chemically competent cells. Two hundred nanograms of plasmid containing the mini-IStron were then delivered by transformation. After recovery (37^°^C for 2 h), transformed cells were spun down at 4,000 g for 5 min and resuspended in 30 µl of LB. Cells were then serially diluted (10x) and plated on LB-agar media containing spectinomycin (100 µg ml^-1^), chloramphenicol (25 µg ml^-1^) and IPTG (0.1 mM), and grown for 16 h at 37^°^C. Plates were imaged in an Amersham Imager 600.

For library DNA cleavage assays, *E. coli* strain BL21(DE3) were transformed with a pEffector plasmid containing a codon optimized *Cbo*TnpB on a pACYC backbone. Single colony isolates were selected to prepare electrocompetent cells. One hundred ng of the pooled IStron library plasmids were mixed with 100 µl of electrocompetent cells and transformed in four separate reactions. After recovery (37^°^C for 2 h), transformed cells were plated on large 245 mm x 245 mm bioassay plates containing LB-agar with spectinomycin (100 µg ml^-1^), chloramphenicol (25 µg ml^-1^) and IPTG (0.1 mM), and grown for 16 h at 37^°^C. Plates were scraped to collect cells at a 350x coverage, and plasmid DNA was isolated using the Qiagen Plasmid Midi Kit.

### IStron TnpA mediated excision assays

In individual IStron variant experiments, chemically competent cells were first transformed with a plasmid encoding tnpA under an inducible lac promoter, and transformants were isolated by plating on LB-agar plates with kanamycin (50 µg ml^-1^). Single colony isolates were selected to prepare chemically competent cells. One hundred nanograms of plasmid containing the mini-IStron were then delivered by transformation. After recovery (37^°^C for 2 h), transformed cells were plated on LB-agar plates containing spectinomycin (100 µg ml^-1^), kanamycin (50 µg ml^-1^) and IPTG (0.5 mM). Following overnight growth at 37^°^C, colonies were scraped and resuspended in LB medium. To prepare cell lysates, approximately 3.2 × 10^8^ cells (equivalent to 200 µl of culture at OD_600_ = 2.0) were transferred to a 96-well plate. Cells were pelleted by centrifugation at 4000 g for 5 min and resuspended in 80 µl of Mili-Q H_2_O. Next, cells were lysed by incubating at 95°C for 10 min in a thermal cycler. The cell debris was pelleted by centrifugation at 4000 g for 5 min, and 10 µl of lysate supernatant was removed and serially diluted in Mili-Q H_2_O to generate 10-fold lysate dilutions for PCR analysis. PCR reactions were performed using primers annealing in 5’ and 3’ IStron-flanking sequences, which can be used to amplify both unexcised loci (longer amplicon) and excised loci (shorter amplicon). Each 20 µl PCR reaction contained 1× OneTaq Master Mix (NEB), 0.2mM of each primer, and 1 µl of 10-fold diluted lysate. Thermal cycling conditions included DNA denaturation (94°C for 30 s), 30 cycles of amplification (denaturation: 94°C for 15 s; annealing: 46°C for 15 s; extension: 68°C for 15 s), followed by a final extension (68°C for 5 min). Products were resolved by 1.5% agarose gel electrophoresis and visualized by staining with SYBR Safe (Thermo Fisher Scientific).

For library TnpA mediated excision assays, *E. coli* strain BL21(DE3) were transformed with a pEffector plasmid containing a codon optimized *Cbo*TnpA on a pCOLA backbone. Single colony isolates were selected to prepare electrocompetent cells. One hundred ng of the pooled IStron library plasmids were mixed with 100 µl of electrocompetent cells and transformed in two separate reactions. After recovery (37^°^C for 2 h), transformed cells were plated on large 245 mm x 245 mm bioassay plates containing LB-agar with spectinomycin (100 µg ml^-1^), kanamycin (50 µg ml^-1^) and IPTG (0.5 mM), and grown for 16 h at 37^°^C. Plates were scraped to collect cells at a 100x coverage, and plasmid DNA was isolated using the Qiagen Plasmid Midi Kit.

### IStron splicing assays

In individual IStron variant experiments, *E. coli* BL21(DE3) strains were transformed with a Mini-IStron variant encoding plasmid. Single colonies were picked from a plate and used to inoculate cultures in LB with spectinomycin (100 µg ml^-1^). Following overnight growth, the cultures were inoculated at 40x dilution in LB supplemented with spectinomycin (100 µg ml^-1^) and IPTG (0.5 mM) and grown until an OD_600_ of 0.5-0.7. An aliquot equivalent to 250 µl of cell suspension at OD_600_ = 0.5 was taken from each culture and centrifuged at 4,000 g for 5 min. The cell pellet was resuspended in 750 µl of Trizol (Thermo Fisher Scientific). After incubating 5 min at room temperature, 150 µl of chloroform was added, and tubes were shaken and centrifuged at 12,000 g for 15 min at 4^°^C. The aqueous phase was transferred to a new tube and mixed with an equal volume of absolute ethanol (>96%), followed by RNA purification using the Monarch RNA Cleanup Kit (NEB). Purified RNA was stored at -80^°^C.

For library scale splicing assays, commercial electrocompetent *E. coli* BL21(DE3) cells (Sigma Aldrich) were transformed with one hundred ng of pooled library plasmids, in three reactions. Following 1 h of recovery at 37^°^C, samples were plated on large 245 mm x 245 mm bioassay plates containing LB-agar with spectinomycin (100 µg ml^-1^) and IPTG (0.5 mM), and grown for 6 h at 37^°^C. Plates were scraped to collect cells at a 100x coverage, followed by RNA extraction and reverse transcription.

For experiments with dTnpB, *E. coli* strain BL21(DE3) was transformed with a pEffector plasmid containing a codon optimized catalytically dead *Cbo*TnpB on a pACYC backbone. Single colony isolates were selected to prepare electrocompetent cells. One hundred ng of the pooled IStron library plasmids were mixed with 100 µl of electrocompetent cells and transformed in three separate reactions. The rest of the workflow was the same as above.

For quantitative measurements, experiments were carried out in biological triplicates. qPCR was performed as previously described (10).

### RNA extraction

Cell pellets were resuspended in 750 µl of Trizol (Thermo Fisher Scientific). After incubating 5 min at room temperature, 150 µl of chloroform was added, and tubes were shaken and centrifuged at 12,000 g for 15 min at 4^°^C. The aqueous phase was transferred to a new tube and mixed with an equal volume of absolute ethanol (>96%), followed by RNA purification using the Monarch RNA Cleanup Kit (NEB). Purified RNA was stored at -80^°^C.

### dsDNase treatment and reverse transcription

Two hundred nanograms of the purified total RNA was used as an input for reverse transcription reactions. First, total RNA was treated with 1 µl of dsDNase (Thermo Fisher Scientific) in a 1× dsDNase reaction buffer in the final volume of 10 µl, incubating at 37°C for 20 min. Then 1 µl of 10 mM dNTP, 1 µl of 2 mM oSL12027, and 1 µl of 2 mM oSL13568 were added for gene-specific priming, and samples were heated at 65°C for 5 min. Tubes were then placed directly on ice, followed by the addition of 4 µl of SSIV buffer, 1 µl of 100 mM dithiothreitol (DTT), 1 µl of SUPERase In (Thermo Fisher Scientific), and 1 µl of Super-Script IV Reverse Transcriptase (200 U µl^−1^,Thermo Fisher Scientific); incubation at 53°C for 10 min; and then incubation at 80°C for 10 min. The resulting cDNA was diluted and used for NGS library prep, endpoint PCR and endpoint qPCR.

### Library assay high-throughput sequencing

100 ng of the isolated IStron plasmid DNA or cDNA were used as input for PCR1 amplification with Kapa DNA polymerase (Roche) for 12 cycles, using specific primers that flank the variable RE position and barcode of each plasmid. Samples were cleaned up and eluted in 30 µl EB. 2 µl of the PCR1 product were amplified in 12 cycles for PCR2. Products were resolved by 1.5% agarose gel electrophoresis. For TnpA mediated excision and splicing assays, PCR products were separated between bands corresponding to excised and unexcised or spliced and unspliced products, respectively. Only the excised or spliced band was cut out of the gel and cleaned up (Qiagen PCR purification kit). Libraries were sequenced on an Element AVITI in paired-end mode with 150 cycles per end.

## RESULTS

### Pooled library design for dissecting IStron functional determinants

The IStron right end (RE) harbors overlapping sequence features required for TnpA-mediated DNA excision, TnpB-mediated RNA-guided DNA cleavage, and group I intron-mediated RNA splicing (10) (**Fig. 1A**). At the RNA level, TnpB activity and splicing are mutually exclusive, as splicing severs the ωRNA scaffold from its guide sequence, while ωRNA maturation severs the group I intron and host mRNA transcript. To dissect the structural requirements that govern these competing functions, we employed a library-based approach with high-throughput functional readouts.

We designed a pooled oligonucleotide library containing thousands of variants of the IStron RE sequence to comprehensively map how sequence perturbations affect each function and thus better understand the balance between IStron activities (**Fig. 1B**). The library encompassed both systematic sequence perturbations (truncations and nucleotide substitutions) and targeted mutations designed to disrupt specific molecular functions: mutations predicted to disrupt TnpA binding to the RE DNA, mutations affecting ωRNA scaffold formation, and mutations targeting the 3’ splice site (3’SS) recognition sequences required for group I intron splicing. Each library member was assigned a unique 10-base pair (bp) barcode positioned downstream of the RE sequence (**Materials and Methods**). For excision and splicing assays, successful activity removes the IStron sequence itself from the DNA or RNA, respectively. The barcode, which remains on the backbone, enables identification of which library member underwent the reaction. Additionally, because each barcode uniquely identifies its cognate RE variant, sequencing only the 10-bp barcode, rather than the full RE sequence, is sufficient to quantify the abundance of each library member across all three assays. The RE library was synthesized as single-stranded oligos, cloned into a mini-IStron donor plasmid, and deep sequenced to confirm that all variants were present and normally distributed (**Fig. S1A**).

By subjecting the library to three distinct functional assays, we systematically evaluated each RE variant’s performance across all IStron activities (**Fig. 1C**). For TnpA-mediated excision, we co-expressed the library with a TnpA effector plasmid. Successful excision removes the IStron from the DNA backbone in a scarless manner, and functional and non-functional variants were distinguished by endpoint PCR. Group I intron splicing was assessed similarly but without any additional effector proteins. Successful splicing removes the IStron from RNA transcripts, and spliced and unspliced products were distinguished through RNA extraction followed by RT-PCR. For ωRNA-guided DNA cleavage, library members were co-expressed with a TnpB effector plasmid. We designed the guide portion of the ωRNA to target the genomic *E. coli lacZ* locus, such that functional ωRNA formation leads to DNA cleavage and cell death, enriching for non-functional variants in the surviving population. All PCR products were analyzed by high-throughput sequencing.

To validate our experimental design, we first confirmed that the downstream barcode sequences did not interfere with IStron function. Individual excision, splicing and DNA cleavage assays comparing IStron variants with identical wild-type (WT) RE sequences but different barcodes showed equivalent activity across all three assays (**Fig S1B-D**), establishing that the barcode itself did not perturb WT IStron activity. We then optimized PCR conditions to minimize recombination artifacts between barcodes and their cognate RE sequences, comparing different DNA polymerases, amounts of input DNA, and PCR cycle numbers (**Fig. S1E**). Even under optimal conditions, barcode uncoupling was still observed within our final pool (**Fig. S1F**). To assess whether uncoupling was sequence-biased, we examined the distribution of incorrectly paired RE sequences for each barcode. For any given barcode, the most frequently associated incorrect RE sequence represented a small fraction of the uncoupled reads (**Fig. S1G**), indicating that uncoupling occurred stochastically across library members rather than favoring specific sequences. To further confirm that this stochastic uncoupling did not obscure functional differences, we examined the behavior of internal controls across biological replicates: a single WT RE sequence carried by four independent barcodes and twelve negative control variants (three distinct non-functional RE sequences, each with four unique barcodes). Despite low, stochastic barcode uncoupling (**Materials and Methods**), WT controls clustered tightly together across replicates, as did negative controls, and the two groups were clearly distinguishable from one another (**Fig. S2A**). We thus conclude that this source of noise does not systematically bias our measurements of functional activity.

To assess reproducibility, each library-scale assay was performed in biological replicates. Comparing each library member’s abundance across replicates showed strong correlation for each assay (**Fig. 1D, Fig. S2B**). We quantified the abundance of each library member by calculating its representation within each pool, normalizing to its abundance in the input library to account for initial representation differences, and further normalized to WT controls to quantify how perturbations influenced IStron activity. To focus on functionally meaningful differences, we applied a threshold of log_2_-fold change >2 or <-2 relative to wild-type. Notably, the DNA cleavage assay differs from the excision and splicing assays in its dynamic range: because it is a negative selection assay, WT ωRNA activity results in complete depletion from the surviving population, precluding identification of variants with enhanced activity. As a result, enrichment and depletion have opposite meanings across assays: functional variants are enriched for excision and splicing, but depleted for DNA cleavage. The excision and splicing assays thus detect both increases and decreases in activity relative to wild-type, producing a broader range of log_2_-fold change values. This comprehensive quality assessment across replicates and functional assays established a robust framework for systematically investigating the molecular determinants governing IStron biology.

### Systematic mutagenesis reveals sequence requirements for IStron multifunctionality

To determine the sequence requirements for each IStron activity at single-nucleotide resolution, we began by analyzing systematic truncations and substitutions of the RE. We designed a series of variants in which nucleotides were progressively replaced with their reverse complement, one nucleotide at a time in the 5’-to-3’ direction (**Fig. 2A**). This approach effectively mimics truncation, since nucleotide identity rather than presence is expected to drive function, while also controlling for RE length, as actual deletions could introduce confounding effects from size differences, either on activity in the cell or on amplification biases in downstream reactions. By measuring how each truncation affected excision, splicing, and DNA cleavage activities, we could precisely map the minimal RE length required for each function. For transposase-mediated DNA excision, we found that only the terminal 39 nucleotides (nt) of the RE were essential, with progressive truncational mutations in upstream sequences having no effect on TnpA-mediated transposition (**Fig. 2B**). Interestingly, splicing showed even less stringent sequence requirements, requiring only the terminal 18 nt of the RE for group I intron activity (**Fig. 2C**). In contrast, DNA cleavage activity required nearly the entire RE sequence, with the removal of 174 nt or more leading to loss of activity (**Fig. 2D**), consistent with the extensive ωRNA structure needed for TnpB-mediated DNA cleavage (10, 20–23). These findings agree with and refine our previous truncation analysis findings (10), and highlight that, while most of the RE sequence is required to maintain ωRNA function, excision and splicing require only short segments nested within the longer ωRNA scaffold region.

**Figure 2.**
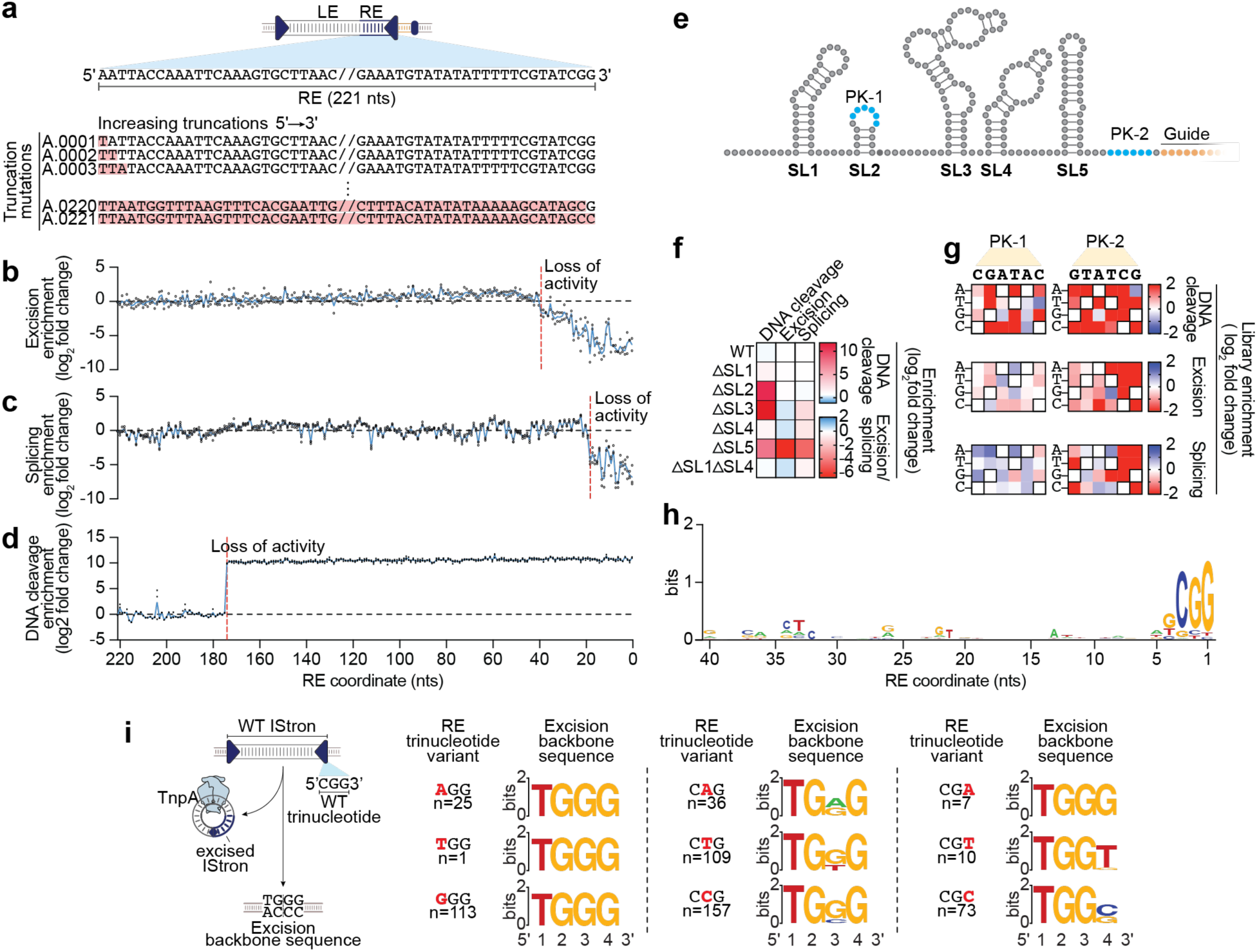
Nucleotide-resolution mapping reveals essential sequence determinants for IStron functions. **a)** Schematic of systematic single-nt truncation analysis across the IStron right end. Truncations were generated by progressively mutating each position to its reverse complement in the 5’-to-3’ direction. **b-d)** Library fold-change patterns for DNA excision (b), RNA splicing (c), and DNA cleavage (d) assays across the RE sequence window, with loss-of-activity positions clearly demarcated (red dashed line). The x-axis shows nucleotide positioning across the RE, 5’ to 3’. Each dot represents a biological replicate, and the blue line connects the mean of these replicates. RE coordinates are defined relative to the IStron RE 3’ boundary. **e)** Schematic of IStron ωRNA secondary structure. **f)** Effect of systematic stem-loop deletions across assays. Heatmaps show library changes for variants with individual or combinatorial deletions of stem-loops SL1-SL5. **g)** Single-nt substitution heatmaps for the PK-1 (left) and PK-2 (right) pseudoknot-forming regions in DNA cleavage (top), excision (middle), and splicing (bottom) assays. Log_2_-fold change values are normalized to wild-type. WT nucleotide identities contain black borders. **h)** Sequence logo revealing consensus motifs required for TnpA-mediated DNA excision. Logo was generated using WebLogo, with each library variant represented proportionally to its abundance in the excision-proficient pool. **i)** Schematic of TnpA-mediated IStron excision showing the relationship between the RE trinucleotide and the resulting backbone sequence (left). Sequence logos depicting the post-excision backbone sequence for library members harboring the indicated RE trinucleotide substitutions (right). Substituted positions are bolded in red; n indicates the number of excision-proficient reads for each variant. The x-axis shows backbone coordinates.

Having established the minimal length requirements, we next used our substitution dataset, in which every nucleotide in the RE was individually changed to every other nucleotide (**Fig. S3A**), to identify critical residues within each functional region. We focused first on DNA cleavage, as it encompasses the largest portion of the RE. A functional ωRNA depends on a specific predicted secondary structure comprising five stem-loops (SL1-SL5) and a pseudoknot (PK) interaction (**Fig. 2E**). While some stem-loop substitutions abolished DNA cleavage activity, the pattern was insufficient to determine which stem-loops were essential (**Fig. S3B**). To systematically assess the contribution of each structural element, we designed targeted deletions of individual stem-loops. Deletion of SL2, SL3 and SL5 abolished DNA cleavage, while deletions of SL1 and SL4 had no effect (**Fig. 2F**). Combinatorial deletions of multiple stem-loops produced no additional phenotype beyond the single deletions (**Fig. S3C**), indicating that loss of any single critical stem-loop was sufficient to inactivate the ωRNA. Of these stem-loops, only SL5 overlaps with the shorter sequences required for excision and splicing, and its deletion accordingly affected all three functions.

Our previous work (10) demonstrated that the pseudoknot is critical for TnpB-mediated DNA cleavage but dispensable for splicing. The pseudoknot is made up of two portions: PK-1, found in SL2 of the ωRNA structure, in a portion that is not required for excision or splicing, and PK-2, found in the 3’ portion of the ωRNA, in nucleotides that are also required for excision and splicing. In agreement with our previous work, nucleotide substitutions in either PK-1 or PK-2 disrupted DNA cleavage activity. However, only substitutions in PK-2 affected excision and splicing, while PK-1 substitutions had no effect, demonstrating that PK-2 represents an overlapping functional requirement for both ωRNA structure, excision, and splice site recognition (**Fig. 2G**).

We next focused on excision, for which previous work has shown that IS607 elements contain short directly-repeated motifs at their ends required for TnpA recognition and binding(24–26). To systematically map critical residues across the full 39 nt excision-competent region, we first visualized our substitution data as a heatmap (**Fig. S3D**), reasoning that substitutions disrupting TnpA binding would reduce excision efficiency and thereby reveal discrete recognition motifs. Although this analysis did not reveal obvious binding sites, we next applied an alternative approach in which each substitution was assigned a weighted score based on its effect on excision activity compared to wild-type, and these scores were then converted into a sequence logo to highlight residues uniquely sensitive to alteration. This analysis similarly indicated that excision sensitivity is broadly distributed across the IStron RE, rather than concentrating into discrete internal motifs (**Fig. 2H**). Because single-nt substitutions can introduce substantial noise, we further averaged substitution effects across sliding windows of 2 nt (**Fig. S3E**) or 4 nt (**Fig. S3F**) to better distinguish regions essential for excision. These analyses confirmed the minimal RE length required for all three activities (**Fig. S3G**) but again suggested the absence of extended TnpA recognition motifs within the element. Instead, our analyses revealed strict sequence requirements at the 3’ terminus. Although the IStron transposase is a small serine recombinase, terminal dinucleotide requirements are best characterized in large serine recombinases, where they determine target-site selectivity and can be altered to redirect integration provided both ends match (27–30). IStrons appear more constrained: we previously showed that most dinucleotide substitutions abolish transposition even when changed at both ends (10). Our sequence logo analysis extends this finding by identifying a critical third nucleotide, indicating that for the IStron RE, a CGG trinucleotide, rather than just a dinucleotide, is required for efficient excision to occur.

To elucidate the role of the terminal RE trinucleotide in excision, we examined whether variations in this sequence affected the excision product itself. In our excision assays, the deep-sequenced product is the excised plasmid backbone containing each variant’s unique 10-nt barcode (**Fig. 1C**), which for the WT IStron, harbors the canonical scarless donor joint sequence, TGGG. All substitutions within the terminal three nucleotides of the RE reduced excision efficiency (**Fig. S3D**). Critically, however, most substitutions to the two most 3’ nucleotides, which compromise the canonical recombinase dinucleotide, altered the sequence of the donor joint, in a substitution-dependent manner (**Fig. 2I**). The only exception, the CGA variant, retained the WT "TGGG" joint. The only exception, the CGA variant, retained the WT TGGG joint, albeit with very few backbone reads (n=7) indicative of a severe excision defect. In contrast, changes to the most 5’ nucleotide of the trinucleotide reduced excision efficiency without affecting the donor joint sequence, which maintained the WT “TGGG” sequence. This pattern is consistent with the conserved chemistry of serine recombinases, in which the two most 3’ nucleotides are cleaved and rejoined during excision, such that a substitution at either position is carried directly into the donor joint. These findings reveal that the terminal trinucleotide serves as a critical convergence point for all three IStron functions. The terminal trinucleotide not only determines excision efficiency and establishes the target sequence for TnpB-mediated DNA cleavage at the donor joint, but also forms part of the group I intron structure required for splicing activity; any variation to these residues severely compromises splicing (**Fig. S3H**). Thus, this short sequence motif represents a remarkable example of molecular multifunctionality, where the same nucleotides are simultaneously constrained by the requirements of DNA transposition, RNA-guided DNA cleavage, and RNA splicing.

### Population level assay comparisons

Our parallel screening approach enabled direct comparison of sequence requirements across the three IStron molecular pathways. To systematically dissect the molecular determinants governing IStron function and RNA fate, we compared the performance of each library member across assays (**Fig. 3A-C**). Comparison of excision and splicing activities (**Fig. 3A**) revealed that most variants maintained WT-like levels of activity (a log_2_-fold change between -2 and 2) in both assays (**Fig. S4A**). Rare variants exhibited enhanced activity relative to wild-type (log_2_-fold change > 2), with some showing an enhanced increase in activity in both excision and splicing simultaneously. Intriguingly, these hyperactive variants (**Fig. S4B**) harbored mutations far upstream of the minimal sequences required for either function, within regions essential for DNA cleavage activity. All library variants exhibiting enhanced splicing activity were completely inactive for TnpB-mediated DNA cleavage, suggesting that ωRNA destabilization can paradoxically enhance IStron splicing.

**Figure 3.**
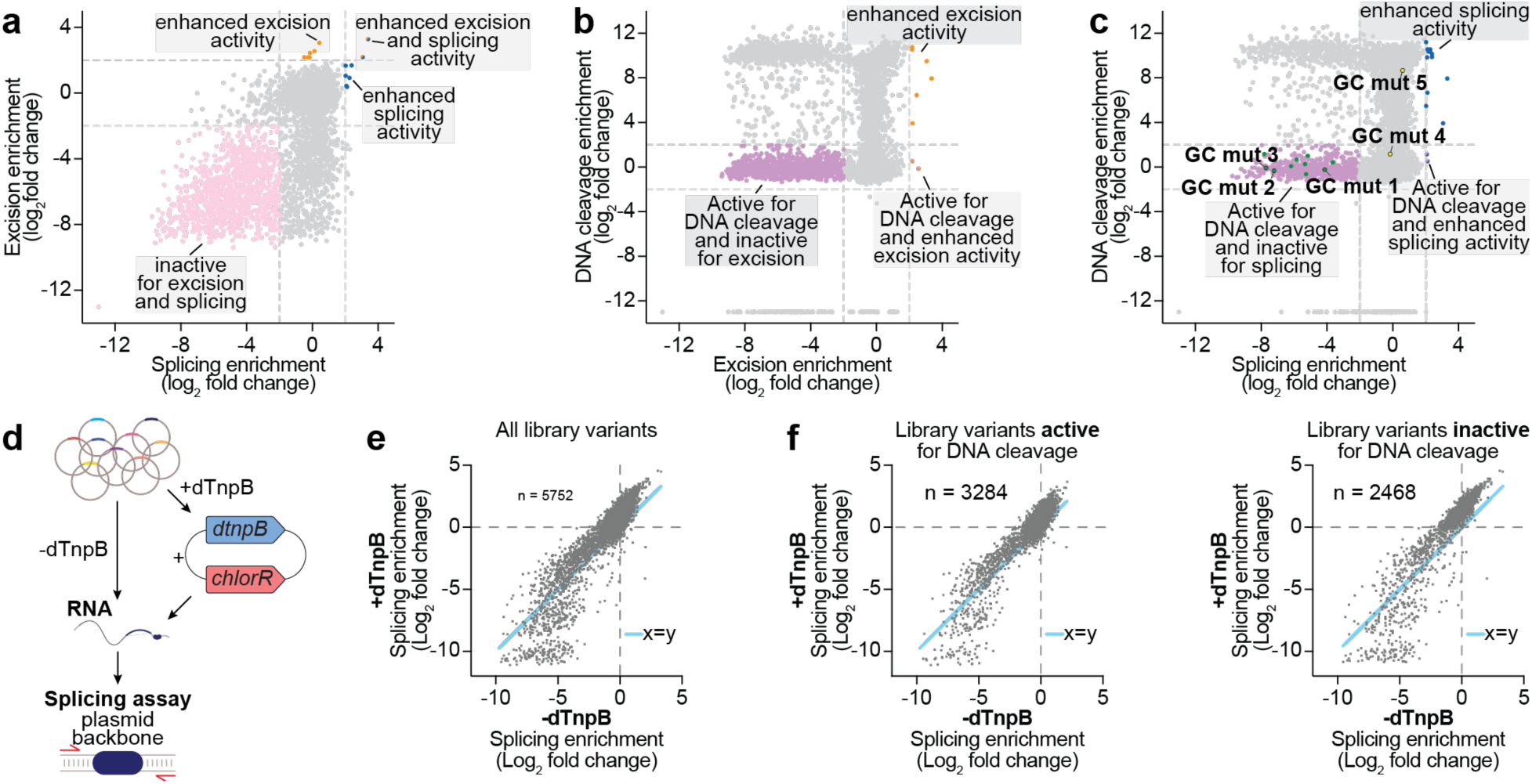
Population level comparisons between library assays. **a)** Direct comparison of library member performance in excision versus splicing assays. Members that were strongly enriched in the excision and splicing library pool (log2-fold-change > 2) were demarcated as ‘enhanced’ and are colored orange and blue, respectively. Members active for more than one function are colored in two colors. Members that were strongly depleted in the library pools (log2 fold-change < -2) were demarcated as ‘inactive’ and colored pink. Log2-fold change of 2 and -2 are marked with a dotted grey line on both axes. Variants absent from the final assay pool were given a log2-fold change value of -13, for visualization purposes. **b)** DNA cleavage versus excision. Data are plotted and colored as in (a). Members that were active for DNA cleavage (log2-fold change <2) are colored purple. **c)** DNA cleavage versus splicing. Data are plotted and colored as in (a) and (b). GC mutants that are active for DNA cleavage but not for splicing are colored green. Representative GC mutants within the ωRNA SL3 are colored in yellow. **d)** Schematic of splicing assay, with or without dTnpB. **e)** Scatter plot of library member enrichment in splicing assay, in the presence (+) or absence (−) of catalytically inactive TnpB (dTnpB) (R² = 0.98). The x=y line shown in blue illustrates the similarity in abundance for each library member with or without dTnpB. Data are plotted for all library variants. **f)** Scatter plot of library member enrichment in splicing assay, in the presence (+) or absence (−) of catalytically inactive TnpB (dTnpB), separated for member that are active for DNA cleavage (left; R² = 0.85) or inactive for DNA cleavage (right; R² = 0.85). Data are plotted as in (e).

We next compared DNA cleavage activity to both excision activity (**Fig. 3B**) and splicing activity (**Fig. 3C**). Both comparisons revealed that library variants segregated into discrete functional populations rather than exhibiting a continuous distribution (**Fig. 3B,C, S4C,D**). Approximately half of the variants retained WT-like activity for both DNA cleavage and excision or splicing, demonstrating that many mutations were tolerated across all three pathways. However, when DNA cleavage was inactivated, variants were roughly twice as likely to retain excision or splicing activity as to lose both functions. This asymmetry reflects the nested architecture of IStron functional elements: because DNA cleavage requires additional sequence elements beyond those needed for excision and splicing, mutations in these extended regions selectively abolish DNA cleavage while preserving the other two functions. Conversely, when excision or splicing is inactivated, DNA cleavage is equally likely to be retained or lost, indicating that the relationship is not strictly nested: some sequence elements within the shared region are specifically required for excision or splicing but dispensable for DNA cleavage.

While these mutational analyses reveal how sequence changes differentially affect IStron pathways, they reflect population-level measurements in which individual transcripts can adopt distinct fates. At the single-molecule level, a more fundamental conflict exists: for any individual RNA transcript, ωRNA maturation and group I intron splicing are mutually exclusive outcomes. Splicing severs the ωRNA guide sequence from its structural scaffold, while ωRNA biogenesis truncates the group I intron structure required for catalysis (**Fig. 1A**). Before investigating the molecular basis for the mutual exclusivity, we first sought to rule out a role for the TnpB protein itself in regulating splicing. We compared library member abundance in the splicing with and without catalytically inactive TnpB (dTnpB) expressed in trans (**Fig. 3D**). Enrichment patterns were indistinguishable between conditions (**Fig. 3E**), and this held true even when we analyzed variants active or inactive for DNA cleavage separately (**Fig. 3F**). These results were unexpected, given our previous finding that WT IStron exhibited reduced splicing activity in the presence of either active or catalytically dead TnpB in individual splicing assays, an effect abolished when the ωRNA scaffold required for TnpB binding was removed (10). The discrepancy likely reflects the greater sensitivity of individual assays to detect modest effects that are masked by noise inherent in library-scale measurements. Regardless, these findings established that TnpB does not substantially influence splicing at the population level, enabling direct comparison of splicing and DNA cleavage assay data to reveal intrinsic RNA-level determinants of fate.

### ωRNA stability determines RNA fate

We hypothesized that increased ωRNA structural stability would favor DNA cleavage over splicing by stabilizing the RNA conformation required for TnpB activity. To test this, we designed a panel of constructs that systematically varied GC base pairing within the ωRNA stem-loops, reasoning that stronger base pairing would stabilize the ωRNA fold. When comparing library performance across DNA cleavage and splicing assays, we found that increased ωRNA stability did not universally inhibit splicing. Only when GC content was increased in SL5 did splicing activity decrease; equivalent increases in other stem-loops such as SL3 had no effect (**Fig. 3C, 4A**). This specificity was intriguing because SL5 is not the only essential stem-loop in the ωRNA secondary structure (**Fig. 2F**), but it is the only one that overlaps with sequences required for group I intron structure formation. We validated these results by individually testing these library variants as cloned mutants in small-scale DNA cleavage and splicing assays (**Fig. 4B**), confirming that increasing the GC content of SL5 by two or more GC base pairs was sufficient to abolish splicing activity while maintaining DNA cleavage activity.

**Figure 4.**
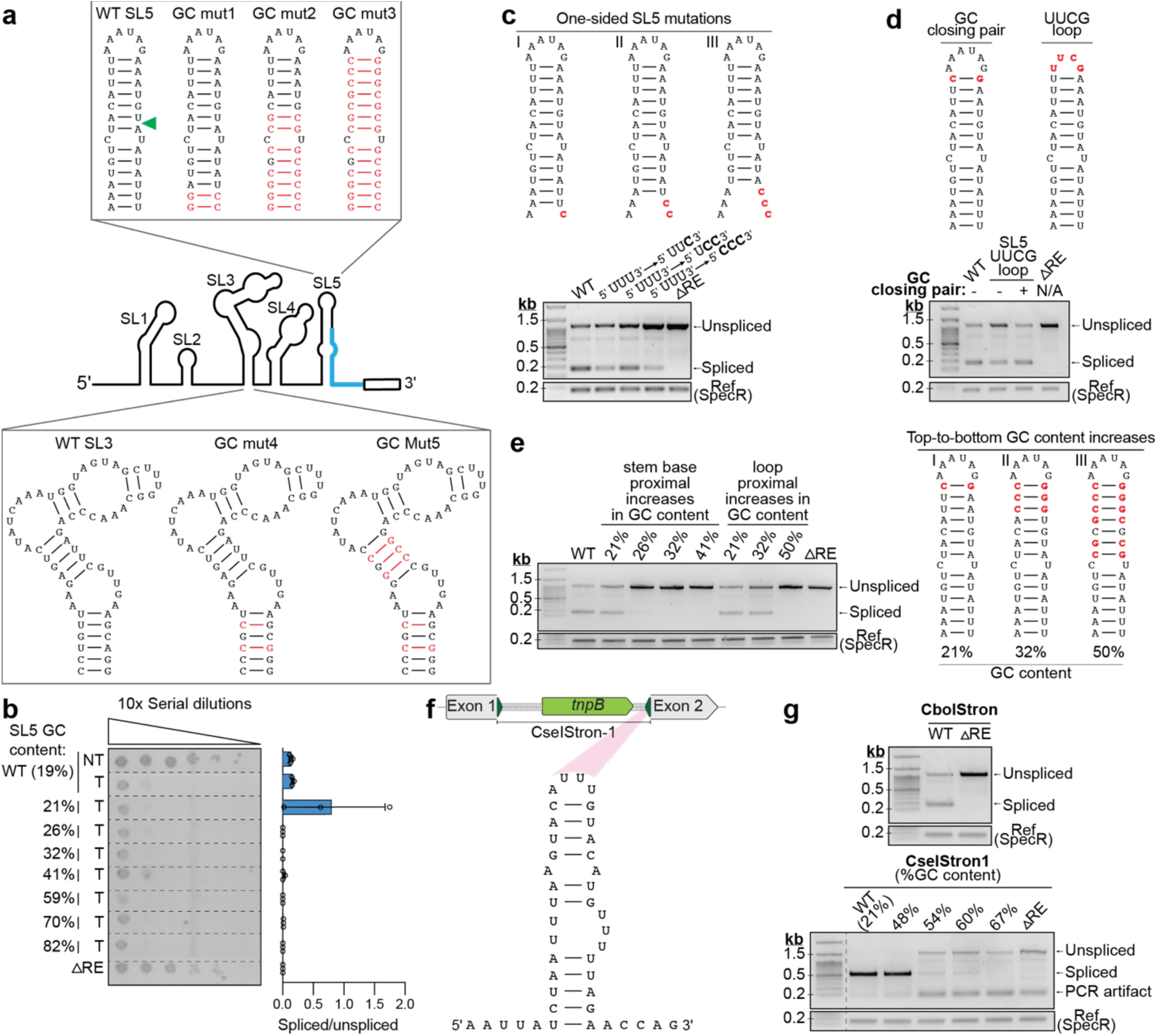
ωRNA structural stability determines RNA fate between splicing and DNA cleavage. **a)** Schematic of WT SL5 and mutants with progressively increasing GC content (GC mut1-3), and equivalent mutations in SL3 (GC mut 4-5). The green triangle shows where the minimal splicing determinant sequence begins; the blue portion of the ωRNA schematic shows the minimal splicing determinant sequence region. **b)** Quantification of DNA cleavage activity (left) and intron splicing activity (right) for ωRNA mutants containing increasing GC richness in SL5. Left, bacterial spot assay demonstrating that all SL5 GC mutants retain RNA-guided DNA cleavage activity. Cells expressing TnpB from a synthetic expression plasmid were transformed with IStron variants containing the indicated SL5 GC content, and 10-fold serial dilutions are shown. Right, RT-qPCR quantification of splicing efficiency for the same IStron variants in the absence of TnpB. RT-qPCR data are means ± SD (n=3). ΔRE, IStron variant containing a scrambled RE sequence. **c)** Effects of one-sided SL5 mutations on splicing. Left, schematic of mutations affecting only the 3’ strand of SL5. Right, agarose gel electrophoresis of RT-PCR products showing unspliced and spliced products (top) relative to a SpecR reference amplicon. ΔRE, IStron variant containing a scrambled RE sequence. **d)** Effects of a stabilizing GC loop-closing pair and UUCG tetraloop substitution in SL5 on splicing. Left, schematic of mutations. Right, data are plotted as in (c). **e)** Effects of loop-proximal (top-to-bottom) increases in GC content compared to base proximal (bottom-to-top) increases in GC content on splicing. Left, schematic of mutations. Right, data are plotted as in (c). **f)** Schematic of *Cse*IStron-1 from *Clostridium senegalense* and the RNA secondary structure of its 3’ most stem loop within the predicted ωRNA structure. **g)** Agarose gel electrophoresis of RT-PCR products from the *Cse*IStron-1 splicing assay, showing unspliced and spliced products for both *Cse*IStron-1 GC mutants (bottom gel) and *Cbo*IStron WT control (top gel), relative to a SpecR reference amplicon. ΔRE, IStron variant containing a scrambled RE sequence for the indicated element.

The overlap between SL5 and group I intron structural elements raised the question of whether the splicing defect arose from altered nucleotide identity in this region or from increased base-pairing stability. To distinguish between these possibilities, we designed one-sided mutations that increased GC content on only one strand of the SL5 stem, thereby changing the nucleotide sequence in the region required for splicing without increasing SL5 stability. These variants retained splicing activity (**Fig. 4C**), indicating that the propensity of SL5 to form a stable stem-loop structure, rather than its primary nucleotide sequence alone, determines the splicing outcome.

Having established that SL5 stability governs RNA fate, we next asked which region of the stem-loop was responsible. We first tested whether the loop sequence itself was a major determinant, by substituting the native loop sequence with a highly stable UUCG tetraloop [(31–34), or by strengthening the loop-closing base pair to a more stable GC pair. Both modifications are expected to increase the thermodynamic stability of SL5, and tetraloop substitution has separately been shown to increase DNA cleavage activity in TnpB systems (21). However, neither modification led to a loss of splicing in the IStron (**Fig. 4D**). We next tested whether the position of stabilizing base pairs within the stem mattered, by designing variants in which the GC content was increased from the loop towards the base of the stem (**Fig. 4E**), rather than at the base of the stem loop, as in our original panel (**Fig. 4B**). This loop-proximal increase in GC content required >50% GC content to abolish splicing, compared to only 26% when GC base pairs were added at the stem base. Loop sequence, loop-closing base pair and loop-proximal increases in GC content had no discernible effects on DNA cleavage activity (**Fig. S5A,B**). Together, these results demonstrate that stability specifically at the base of SL5 governs RNA fate.

In bacteria, RNA folds co-transcriptionally, and the rate of transcription elongation, along with polymerase pausing, can direct a nascent transcript down alternative folding pathways (35–37). We therefore asked whether the rate of IStron transcription influences RNA fate. We constructed strains carrying an *rpoB* mutation (H447P) that slows RNA polymerase and delays synthesis of the full-length transcript (38), and compared IStron activity to a wild-type strain. We observed no significant difference in DNA cleavage or splicing relative to WT (**Fig. S5C**), indicating that transcription kinetics do not detectably contribute to RNA fate.

Finally, we sought to determine whether this relationship between splicing and stability at the base of the stem loop is conserved across IStrons. We previously identified a related IS607-family IStron from *Clostridium senegalense*, *Cse*IStron-1, which we have shown exhibits robust splicing activity (10). Using an ωRNA covariance model, we predicted the 3’ most stem loop structure (**Fig 4F**) and systematically increased its GC content starting from the base of the stem, as in our original panel with each mutation increasing the GC content. This progressive stabilization resulted in loss of splicing activity in CseIStron-1 (**Fig. 4G**), mirroring our observations with CboIStron. These experiments establish that stability at the base of the 3’-most ωRNA stem-loop is a conserved determinant of RNA fate.

### IStron 3’ splice site selection determinants and alternative splicing

Group I intron RNAs catalyze their own excision through formation of complex, highly specific secondary structures (39–42) that then fold into a tertiary structure with a conserved common core (43–46). This structure comprises a specific arrangement of stem loops connected by junctions, terminating in a 3’ conserved guanosine, the omega G (ΩG), which plays a role in 3’ splice site (3’SS) recognition. For IStrons, most of this catalytic architecture is encoded within the element’s LE, with a small but critical structural contribution from the RE that completes the secondary structure and includes the ΩG (**Fig. 5A**). Accurate splicing is essential for group I introns; splicing even a few nucleotides upstream or downstream of the intron boundaries introduces insertions or deletions that render the spliced transcript non-functional. We therefore sought to determine the molecular features governing 3’SS selection and the factors that influence splicing precision.

**Figure 5.**
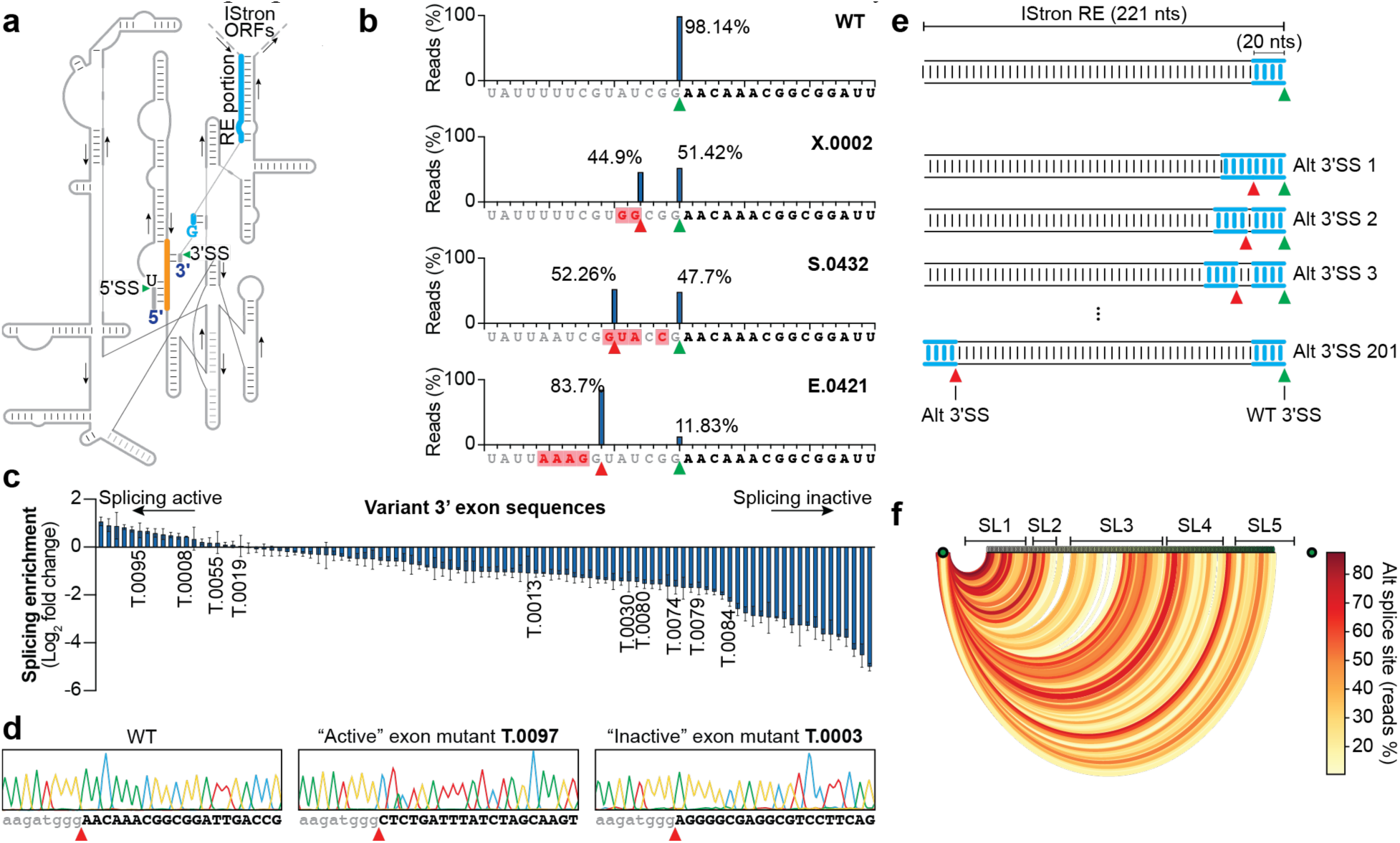
Determinants of precise 3’ splice site selection in IStrons. **a)** Schematic of IStron group I intron structure. Green triangles denote 5’ and 3’ splice sites. Thick blue lines indicate the region of group I intron structure located within the RE. Orange denotes the region within the group I intron where the 3’ exon is expected to base pair. The terminal G nucleotide required for splicing is highlighted. **b)** Distribution of reads from library-scale splicing experiments, showing the exact 3′ splice site used by each library member. Splice sites were classified as WT (spliced at the accurate IStron boundary) or alternative. A representative WT member and three members with varying degrees of alternative splice site usage are shown (X.0002: 44.91%, S.0432: 52.26%, E.0421: 83.7%). Library variant accessions are at the top right of the graph; nucleotide sequences are shown below each distribution; grey lettering denotes IStron RE sequence; bold lettering denotes the 3’ exon; red lettering in red boxes indicates mutations within that member compared to the WT sequence; green triangle denotes WT splice site; red triangle denotes alternative splice site. **c)** Enrichment of library members with different 3’ exon sequences in splicing assays. Abundances are calculated as log_2_-fold change normalized to wild-type. Accessions are below members that were randomly selected for downstream analyses. **d)** Sanger sequencing of PCR products from individual splicing experiments with 3’ exon variants. The sequence of the spliced transcript is written below; the 3’ exon sequence is in bold; red triangles denote splice sites. **e)** Schematic of experiments where the RE portion required for splicing is duplicated within the RE and shifted in the 5’ direction in 1-nt increments. The WT 3’ splice site is marked with a green arrow; a new potential alternative splice site is marked with a red arrow. **f)** Arch diagram depicting alternative splice site choice for library members described in (e). For each member, reads corresponding to the potential alternative splice site generated in the mutant were counted and displayed as a percentage of total reads. Dark green circles denote the WT 5’ and 3’ splice sites. All arches start at the same 5’ splice site and end at different nucleotide positions in the RE in 1-nt increments. As the additional 3’ splice site moves in the 5’ direction, the corresponding green circle becomes lighter. Arch color represents the percentage of alternative splice site choice over WT splice site, as indicated by the color scale. Above the green circles are the positions of different stem loops of the ωRNA.

We started by analyzing the accuracy of 3’SS selection across our library. As with excision products, the deep-sequenced splicing product is the spliced transcript backbone containing each variant’s unique 10-nt barcode. For WT IStron, this yields the canonical scarless spliced product: 98% of the WT reads mapped to the precise IStron boundary (**Fig. 5B**), with the remaining 2% not mapping to alternative positions within the WT IStron or to any other library member. Strikingly, numerous library members selected alternative splice sites to varying degrees, deviating from the WT 3’SS (**Fig. 5B**). To understand what features influenced splice site choice in these variants, we examined the last 20 nt in the RE directly upstream of the alternative splice site for the 100 variants with highest alternative splice percentages, which varied from 20-100%. This analysis revealed that all alternative splice sites share a terminal G, with the majority exhibiting a terminal CGG trinucleotide, with an AT-rich preference across the remaining upstream sequence (**Fig. S6A**). Notably, alternative splice sites were only selected within a narrow window extending up to 10 nt from the WT 3’SS.

The 3’ exon sequence is known to influence group I intron splicing through base-pairing interactions within the intron structure that guide proper 3’SS recognition (43, 47). Mutating the 3’ exon sequence has been shown to lead to an increased use of a cryptic 3’SS (48). Within our library, 1-nt substitution data revealed that not all nucleotide identities within the 3’ exon sequence were tolerated for splicing (**Fig. S6B**). To systematically assess the contribution of the 3’ exon sequence to splicing efficiency and precision, we tested 100 variants containing the WT RE paired with random 3’ exon sequences harboring varying GC composition; these variants were originally designed as controls for DNA cleavage assays, since altering this sequence disrupts ωRNA targeting of the genome. Surprisingly, the variants displayed a wide range of splicing efficiencies, from WT-like levels to a complete loss of splicing activity (**Fig. 5C**). To validate these results, we selected the five most active and five least active variants for individual splicing assays. Analysis of the 3’ exon sequences of these variants did not reveal a clear sequence determinant underlying the variance in splicing efficiencies (**Fig. S6C**), yet RT-qPCR and RT-PCR confirmed that 3’ exon variation alone was sufficient to substantially reduce splicing activity (**Fig. S6D,E**). However, Sanger sequencing of gel-purified products corresponding to the expected splice product size revealed that all variants, regardless of exon sequence or splicing efficiency, maintained strict fidelity to the WT 3’SS selection at the IStron boundary (**Fig. 5D, S6F**). To corroborate this finding, we examined 3’SS selection across randomly selected library members with diverse 3’ exon sequences (**Fig. 5C**). Most of these variants selected the correct 3’SS, although in some cases the 3’ exon sequence promoted usage of an alternative splice site at lower frequencies than at the IStron boundary (**Fig. S6G**). These findings suggest that the 3’ exon sequence affects the efficiency with which the proper 3’SS is selected more than the accuracy with which it is selected.

Having established that 3’ exon sequence influences the efficiency of 3’SS selection more than its accuracy, we next asked what structural features within the RE promote faithful splice site recognition. The minimal RE sequence required for splicing comprises only 18 nt at the 3’ terminus, with the remainder dedicated to ωRNA structure formation (**Fig 2C-D**). We therefore hypothesized that the extended ωRNA structure upstream of these minimal determinants might sterically restrict 3’SS accessibility, confining splice site selection to the native 3’ boundary. To test whether alternative splice sites could be recognized when placed within the structured ωRNA region, we generated a series of constructs containing two identical copies of the 3’-terminal splicing determinant sequence. Each construct contained one copy at the native 3’ position and a second copy positioned progressively upstream in 1-nt increments, replacing the corresponding internal RE sequence (**Fig. 5E**). Deep sequencing of spliced products from these variants, which present the splicing machinery with a choice between the native and alternative 3’SS, revealed how alternative splice site selection efficiency varied as a function of position within the RE (**Fig. 5F**). If splice site selection were governed primarily by proximity to the LE, which encodes most of the group I intron catalytic structure, alternative splice site usage would be expected to increase as the duplicated site approached the 5’ end of the RE. Instead, alternative splice sites were preferentially selected at some positions but not others, revealing a position-dependent pattern that correlated with ωRNA structural features rather than distance from the LE. Alternative sites that fell within non-essential regions, such as SL1, were preferentially selected over the WT 3’SS, consistent with replacement of a dispensable structural element that does not contribute to steric occlusion. Conversely, alternative splice sites that disrupted essential stem loops SL2, SL3, or SL5, thereby inactivating the ωRNA, were also preferentially selected in some cases. However, dispensability or ωRNA disruption alone was not sufficient for alternative splice site selection, as not all variants meeting these criteria preferentially used the alternative site. This suggests that the newly generated 3’ exon sequence, created by each internal duplication, also influences the efficiency of alternative 3’SS recognition, consistent with the role of 3’ exon sequence in modulating splicing efficiency discussed above. Collectively, these results demonstrate that a stably folded ωRNA structure disfavors use of alternative splice sites embedded within it, constraining 3’SS selection to the native intron–exon boundary, with additional contributions from the downstream exon sequence in determining splicing efficiency at any given site.

## DISCUSSION

IStron elements must balance three mutually exclusive molecular functions — TnpA-mediated excision, TnpB-mediated DNA cleavage, and group I intron-mediated self-splicing — to proliferate within their host genomes without driving themselves to extinction or overly compromising host fitness. Through high-throughput functional screening of thousands of systematically designed IStron RE variants, we mapped the sequence requirements governing each function at nucleotide resolution and revealed how these competing activities are coordinated. Our findings establish that IStron sequences have evolved under opposing selective pressures that create an intrinsic functional trade-off.

Systematic mutagenesis revealed that IStron RE sequences are constrained by overlapping functional requirements across all three pathways. The terminal CGG trinucleotide at the 3’ end of the RE represents a convergence point where all three IStron pathways impose overlapping sequence constraints: any single nucleotide change to this trinucleotide severely compromises all three activities, creating a functional linchpin governing IStron evolution. The differential behavior we observe within this trinucleotide is consistent with the conserved chemistry of serine recombinases, in which DNA cleavage generates 2-nt 3’ overhangs that must base-pair for productive ligation following recombinase subunit rotation (49, 50). The two most 3’ nucleotides in the IStron likely constitute the crossover dinucleotide, explaining why their substitution alters donor joint identity in a sequence-dependent manner, in which the mutant overhang is incorporated directly into the ligation product. By contrast, the 5’ nucleotide of the trinucleotide occupies a position adjacent to the crossover site, where perturbations in other serine recombinase systems reduce recombination efficiency by disrupting conformational changes required for strand exchange (51–53). Our data thus provide systematic evidence that terminal dinucleotide requirements in a small serine recombinase parallel those established for large serine recombinases (27–30), while simultaneously revealing that these same nucleotides are subject to additional selective constraints, from ωRNA pseudoknot formation to splice site recognition, that have no parallel in other characterized recombinase systems.

For any individual IStron transcript, ωRNA maturation and group I intron splicing are mutually exclusive outcomes, and RNA structural determinants alone, independent of TnpB, govern this choice. Multiple lines of evidence establish ωRNA formation as the preferred fate. Increased base-pairing stability at the base of SL5, the sole stem-loop overlapping group I intron structural elements, is sufficient to abolish splicing while maintaining DNA cleavage, and this relationship is conserved in the IS607-family *Cse*IStron-1. Because this stem-loop overlaps the intron’s 3′-exon pairing region (Fig. 5A), the effect likely reflects a local competition in which stabilizing the SL5 base sequesters nucleotides that would otherwise remain available for the intron’s internal guide sequence pairing, rather than a shift in the global folding equilibrium of the transcript. Moreover, a stably folded ωRNA disfavors use of potential alternative splice sites embedded within it, confining 3’SS selection to the native intron–exon boundary. Our previous work suggested that IStron RNA fate might be governed by dynamic, context-dependent features analogous to riboswitch-mediated regulation of ancestral group I introns (10), (54). However, the systematic data presented here reveal a more rigid architecture in which, rather than toggling between functional conformations, IStron RNA fate is primarily dictated by the intrinsic thermodynamic stability of the ωRNA scaffold. This architecture prioritizes transposon maintenance through TnpB-mediated donor joint cleavage and homologous recombination over the host-protective benefit of splicing. *Cse*IStron-1, which exhibits robust splicing but no detectable DNA cleavage activity (10), illustrates the extreme consequence of this rigidity: high splicing efficiency may only be achievable when the ωRNA maintenance function is effectively abandoned. We note that our assays isolate this RNA-intrinsic determinant, as the mini-IStron substrate is not translated across the fate-determining structures; co-translational or other trans-acting effects in the native element could further modulate this balance. More broadly, this suggests that multifunctional noncoding RNAs encoding competing molecular fates may be inherently constrained in their capacity to optimize any single function, and that escaping such constraints requires relinquishing one activity entirely rather than fine-tuning a dynamic equilibrium.

Interestingly, group I intron splicing proved to be remarkably sensitive to the sequence identity of the 3’ exon. Random 3’ exon sequences paired with the WT IStron RE yielded highly variable but reproducible splicing efficiencies, from WT levels to almost complete losses of activity. This variation likely reflects differential base-pairing between the 3’ exon and the core group I intron structure, which is known to guide proper 3’ splice site recognition in other self-splicing introns (43, 47, 48). Throughout the IStron life cycle, TnpA-mediated transposition to new genomic sites generates novel 3’ exon contexts, and a substantial fraction of these may be inherently incompatible with efficient splicing. This external dependence diminishes the host-protective benefit that splicing provides, and may in part explain why ωRNA formation is so heavily favored over splicing: DNA cleavage activity is robust to genomic context, whereas splicing is not. The preservation of splice site accuracy across diverse 3’ exon sequences nonetheless ensures that when splicing does occur, it faithfully restores the interrupted transcript, maintaining at least partial host-protective function even at suboptimal insertion sites.

Several technical limitations shape the interpretation of our findings. First, because the ωRNA occupies most of the RE sequence, the mutations we designed potentially create an ascertainment bias toward ωRNA-disrupting variants. Second, the DNA cleavage assay we employed relied on library member depletion, in which functional ωRNAs led to cell death. While this approach effectively identified loss-of-function mutations, it precluded discovery of hyperactive variants, as wild-type IStrons were already completely depleted from surviving populations. Third, library-scale experiments yielded population-level measurements relative to controls, rather than single-variant characterization. We addressed this through targeted validation of individual clones for select findings, though the scale of the library precluded comprehensive single-variant characterization. Finally, all experiments were performed in E. coli BL21(DE3) using over-expression vectors with synthetic promoters, which constrains the physiological interpretation of our findings. Splicing efficiencies may differ in the native host, though the sequence-intrinsic mechanism governing the balance between ωRNA maturation and splicing should still hold. In addition, tnpA and tnpB are natively encoded within the IStron under a single promoter, whereas our trans-expression system alters their levels and relative stoichiometry.

Several important questions emerge from this work. The apparent hierarchy between DNA cleavage and splicing — suggested by *Cse*IStron-1’s high splicing activity coupled with undetectable DNA cleavage — raises broader questions about the evolutionary trajectories available to IStron elements. Whether distinct IStron lineages across diverse hosts have converged on similar solutions to the splicing–cleavage trade-off, or whether different selective environments favor different positions along this functional spectrum, remains to be determined. IStrons exemplify how mobile genetic elements can evolve sophisticated regulatory mechanisms to manage their own proliferation, balancing selfish replication with the need to avoid host extinction. Understanding these regulatory mechanisms provides insights into the evolutionary dynamics of transposable elements and their long-term persistence in host genomes.

## DATA AVAILABILITY

High-throughput sequencing data are available at the National Center for Biotechnology Information (NCBI) Sequence Read Archive (BioProject Accession: PRJNA1432764). Custom scripts used for analyses of high-throughput sequencing data are available at GitHub (https://github.com/sternberglab/Mortman_et_al_2026). Datasets generated and analyzed in the current study are available from the corresponding authors on reasonable request.

## SUPPLEMENTARY DATA STATEMENT

Supplementary data are available at NAR online.

## ACKNOWLEDGMENTS

We thank R. Žedaveintė for her help in assay design, data interpretation, and support throughout this project; M. Walker for his guidance in library-scale workflows and analysis; C. Meers and G.D. Lampe for meaningful discussion; K.Y. Ralwins de la Rosa, A.I. Palmieri, A.J. Robinson, and T.M. Smith for laboratory support; L.F. Landweber for qPCR instrument access; and the JP Sulzberger Columbia Genome Center for NGS support.

## AUTHOR CONTRIBUTIONS

E.E.M. and S.H.S. conceived the project. E.E.M. performed all experiments and analyses and wrote the manuscript, with input from S.H.S.

## COMPETING INTERESTS

S.H.S. is a co-founder and scientific advisor to Dahlia Biosciences, a scientific advisor to CrisprBits and Prime Medicine, and an equity holder in Dahlia Biosciences and CrisprBits. The rest of the authors declare no competing interests.

Correspondence and requests for materials should be addressed to S.H.S.

