## Supplemental Figures for "Large-scale mutational analysis uncovers molecular mechanisms governing dual RNA functions in transposons"

### SUPPLEMENTARY FIGURES

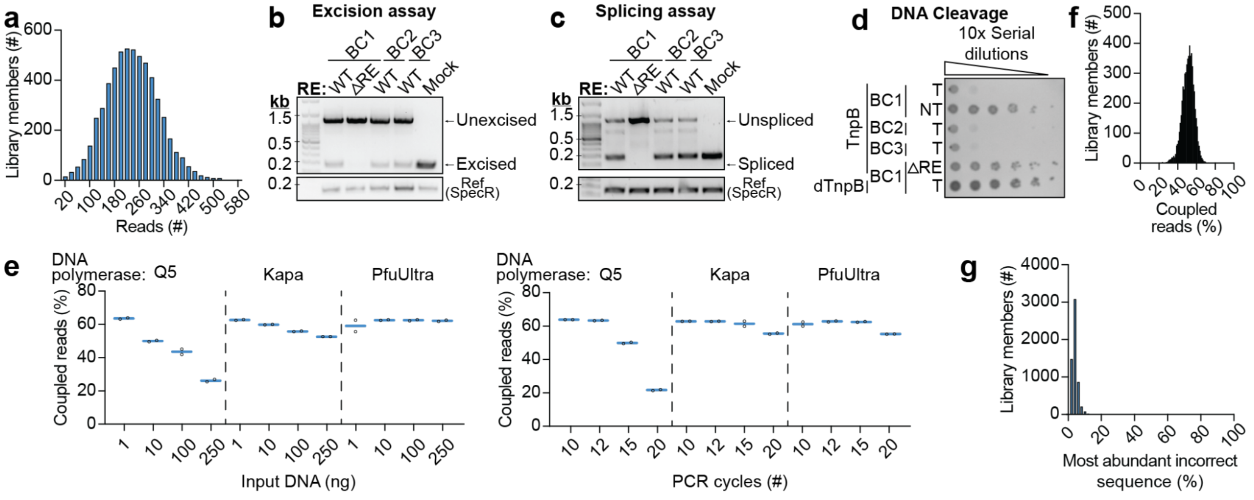


**Figure S1 | Optimization of library preparation for NGS and quality verification.**
**a)** Read count distribution across library barcodes. All library members were present in the final input pool.
**b)** Gel electrophoresis of individual excision assays comparing performance of WT IStron with varying barcode sequences. Cells containing a TnpA expression plasmid were transformed with different IStron donor plasmids. Expected substrates and products of IStron excision generated by PCR are indicated. Mock denotes a positive excision control; the SpecR drug marker was amplified as a positive control for presence of the plasmid.
**c)** Gel electrophoresis of individual splicing assays comparing performance of WT IStron with varying barcode sequences. Cells were transformed with different IStron donor plasmids. RNA was extracted from these cells followed by RT-PCR, the products of which are indicated. Mock and SpecR are as in (b).
**d)** Bacterial spot assays of individual DNA cleavage assays comparing performance of WT IStron with varying barcode sequences. Cells containing a TnpB expressing plasmid were transformed with different IStron donor plasmids with guide portions that had either target-containing (T) or non-target-containing (NT) sequences. Transformants were serially diluted (10x), plated on selective media, and cultured at for 16 h at 37 °C. Additional controls included a mutant with a scrambled RE sequence and a dTnpB control.
**e)** NGS sequencing of the input library to find optimal conditions for amplicon sequencing with minimal barcode uncoupling between variants. To optimize our sequencing output, we tested multiple conditions: the amount of starting input DNA (ng), the number of cycles during the PCR for amplicon sequencing, and the polymerase used. When sequencing the input library prior to performing different assays, the entire library member's RE was sequenced in addition to the barcode, allowing us to analyze each read for whether the barcode was associated with its expected RE (coupled read) or not (uncoupled read). The coupled reads percentage is calculated as the total number of coupled reads divided by the total number of reads for that library, multiplied by 100%.
**f)** Histogram showing the percentage of properly coupled reads for each library member within the entire input pool, calculated as the total number of coupled reads divided by the total number of reads for that barcode.

**g)** Histogram showing the maximum percentage of the most abundant uncoupled read for each library member. For each library member, the most abundant incorrect sequence was identified, and its read count was expressed as a percentage of that member's total reads. These analyses demonstrate that uncoupled reads occur stochastically across library members.

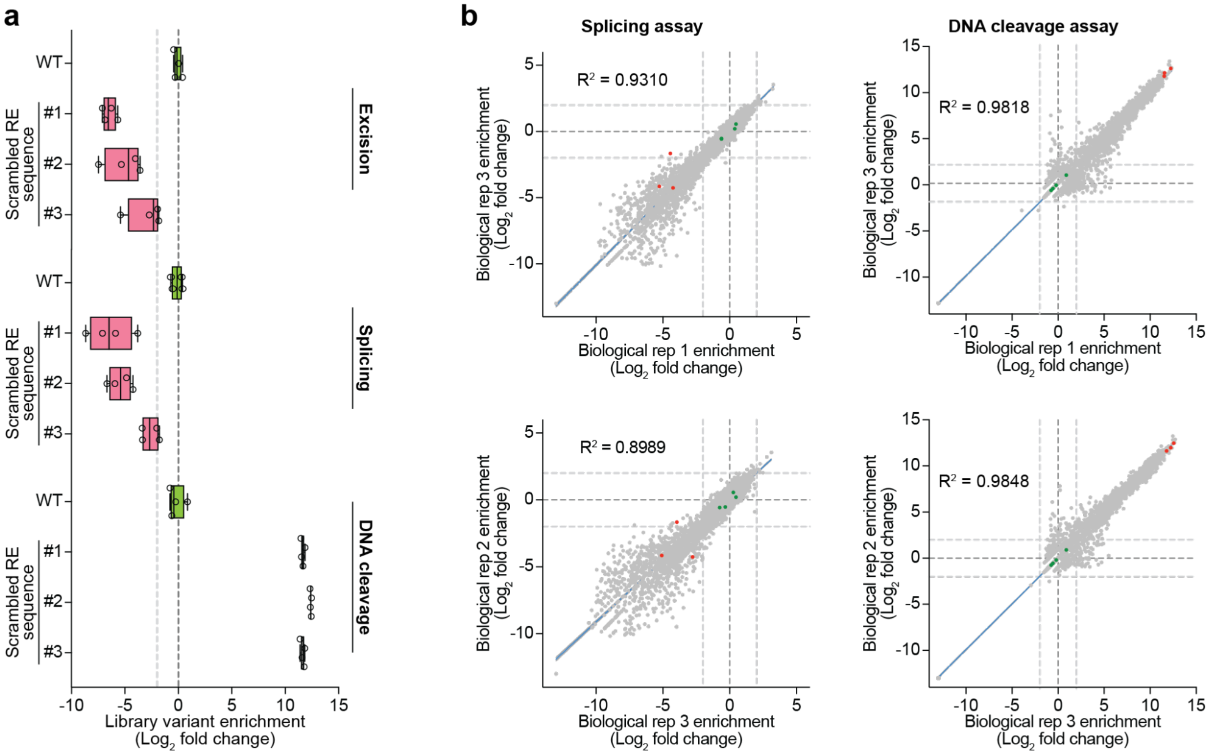


**Figure S2 | Validation of library experimental design.**

**a)** Abundance of WT and negative control members containing scrambled RE sequences. There are four library members for each WT and negative control sequence, containing different barcode sequences. Read counts were normalized to pool size, then to input abundance to calculate fold-enrichment, and finally scaled to WT levels. Data are shown on a $\log_2$ scale.

**b)** Additional biological replicate correlation. Scatter plots show $\log_2$-fold change enrichment of library members normalized to wild-type for splicing (left,) and DNA cleavage (right) assays. $R^2$ values are shown within each graph. WT IStron library members are in green; representatives of each scrambled RE negative control are in red; light gray dotted lines denote the $\log_2$-fold change of 2 and -2.

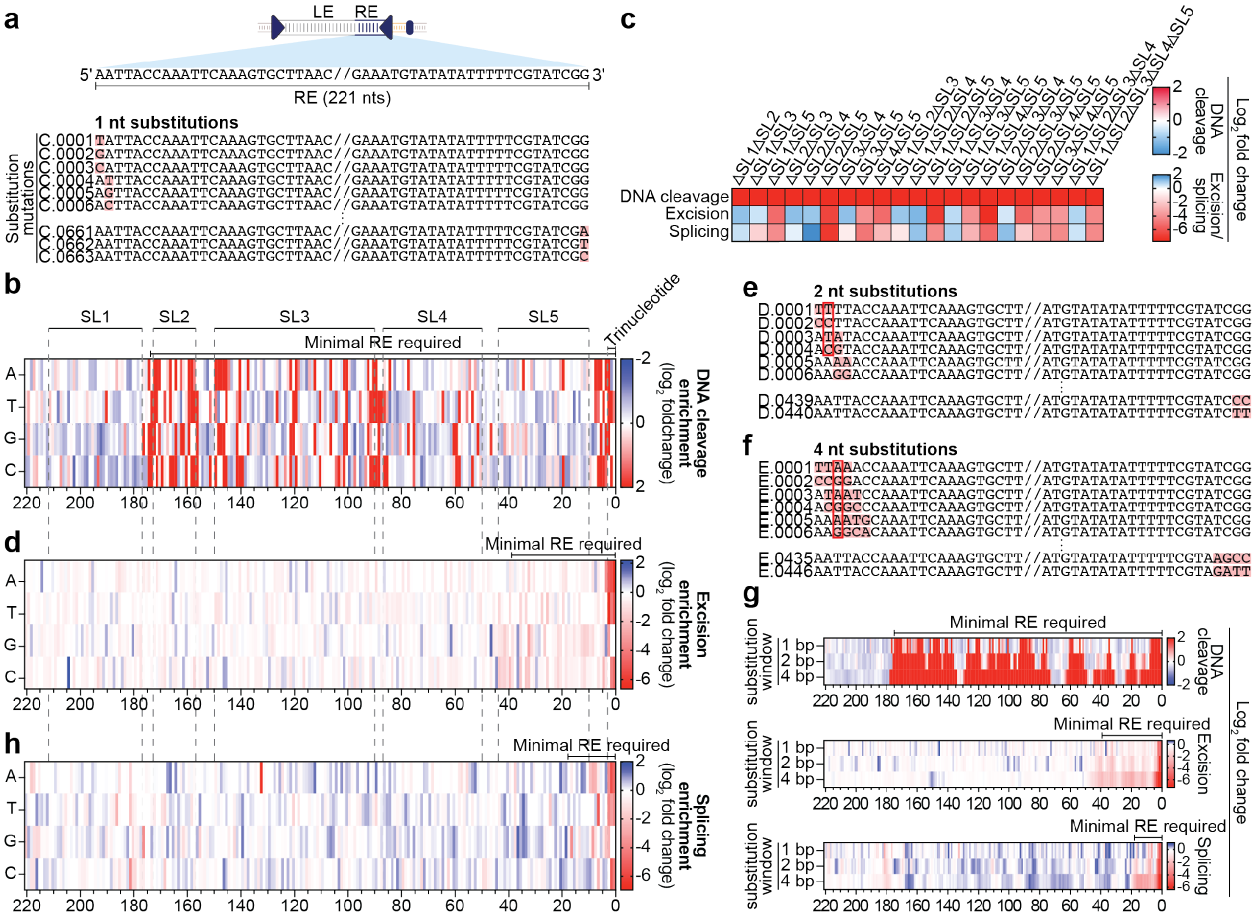


**Figure S3 | Effects of nt substitutions on library members' abundance across assays.**

**a)** Schematic of 1-nt substitutions in IStron RE. The WT IStron is shown above with left end (LE) and right end (RE) boundaries indicated, as well as a magnified view of the RE sequence. Each nucleotide within the RE was substituted to every other possible nucleotide, with pink shading indicating the exact mutation within the library member. Library variant accessions are shown on the left.

**b)** Heatmap depicting $log_2$-fold change in DNA cleavage assay (relative to WT) for each nucleotide substitution across the IStron RE. RE coordinates are on the x-axis. Minimal RE required for this assay indicated above the heatmap. Grey dotted lines indicate the location of the 3' terminal trinucleotide and the ωRNA stem-loops (SL1-SL5). WT nucleotide identities are depicted as white boxes.

**c)** Heatmap depicting $log_2$-fold change (relative to WT) for combinatorial deletions of ωRNA stem-loops in DNA cleavage (top), excision (middle), and splicing (bottom) assays.

**d)** Heatmap depicting $log_2$-fold change in excision assay (relative to WT) for each nucleotide substitution the RE. Data are plotted as in (b).

**e)** Schematic of 2-nt substitutions in the IStron RE. Two consecutive nucleotides were substituted simultaneously, with pink shading indicating mutations. Library variant accessions are shown on the left. Red box highlights an example of library variants whose fold-change values were averaged to derive a single positional value.

**f)** Schematic of 4-nt substitutions in the IStron RE. Four consecutive nucleotides were substituted simultaneously, with pink shading indicating mutations. Library variant accessions are shown on the left. Red box highlights an example of library variants whose fold-change values were averaged to derive a single positional value.

**g)** Heatmaps depicting $log_2$-fold change (relative to WT) for DNA cleavage (top), excision (middle), and splicing (bottom) activities, comparing 1-nt, 2-nt, and 4-nt substitution window libraries. Minimal RE length required for its respective assay indicated above each assay heatmap. For 2-nt and 4-nt libraries, fold-change values were averaged across all library members containing substitutions at each position.

**h)** Heatmap depicting $log_2$-fold change in splicing assay (relative to WT) for each nucleotide substitution the RE. Data are plotted as in (b).

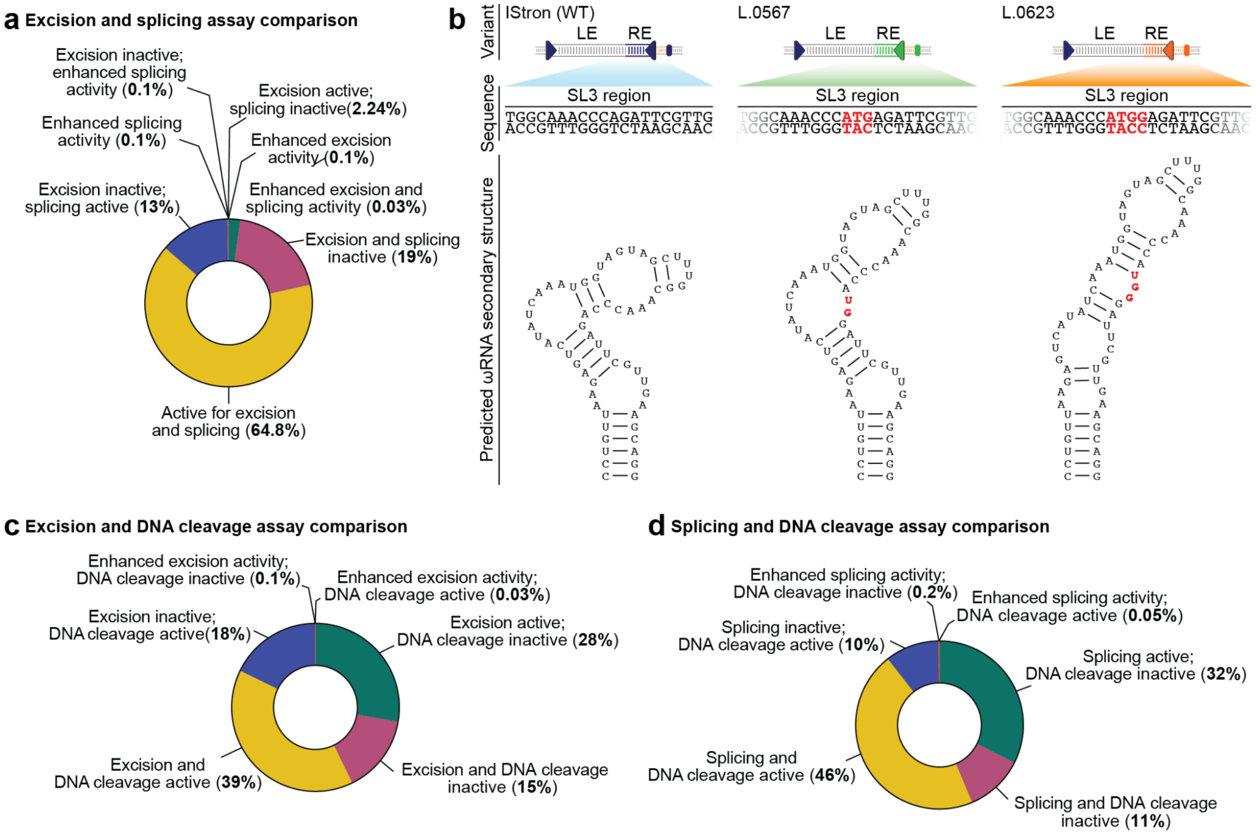


**Figure S4 | Library member categorization across assays.**

**a)** Pie charts depicting the breakdown of library members when comparing between excision and splicing assays. The percentage of library members falling into each category are shown in parentheses, out of 5752 total library members.

**b)** Rare library variants that exhibit enhanced activity in both excision and splicing assays. WT IStron is shown on the left. Variant accessions are shown above. The mutated sequence is highlighted in red. In both cases, these mutations fall within the ωRNA SL3 region. The predicted effect on the ωRNA secondary structure is shown at the bottom.

**c)** Pie charts depicting the breakdown of library members when comparing between excision and DNA cleavage assays. Data are plotted as in (a).

**d)** Pie charts depicting the breakdown of library members when comparing between splicing and DNA cleavage assays. Data are plotted as in (a).

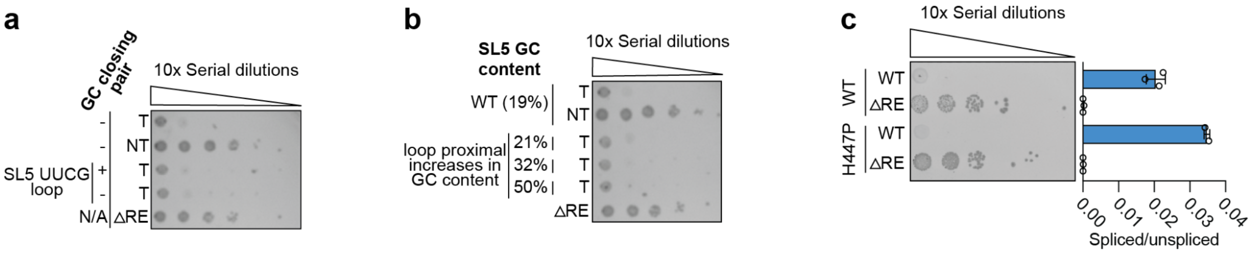


**Figure S5 | SL5 stability modifications do not affect RNA-guided DNA cleavage activity, and slowed transcription in an *rpoB* mutant affects neither DNA cleavage nor splicing.**

**a)** Bacterial spot assays demonstrating the effects of a stabilizing GC loop-closing pair and UUCG tetraloop substitution in SL5 on DNA cleavage activity. Cells expressing TnpB from a synthetic expression plasmid were transformed with IStron variants, and 10-fold serial dilutions are shown.

**b)** Bacterial spot assays demonstrating the effects of loop-proximal (top-to-bottom) increases in GC content in SL5 on DNA cleavage activity. Experiment performed as in (a).

**c)** Quantification of DNA cleavage activity (left) and intron splicing activity (right) for in WT RpoB background and H447P RpoB mutant background. Left, bacterial spot assay demonstrating that RNA-guided DNA cleavage activity is retained in both contexts; Cells expressing TnpB from a synthetic expression plasmid were transformed with IStron variants containing WT or ▵RE IStron, and 10-fold serial dilutions are shown. Right, RT-qPCR quantification of splicing efficiency for the same IStron variants in the absence of TnpB. RT-qPCR data are means ± SD (n=3). ▵RE, IStron variant containing a scrambled RE sequence.

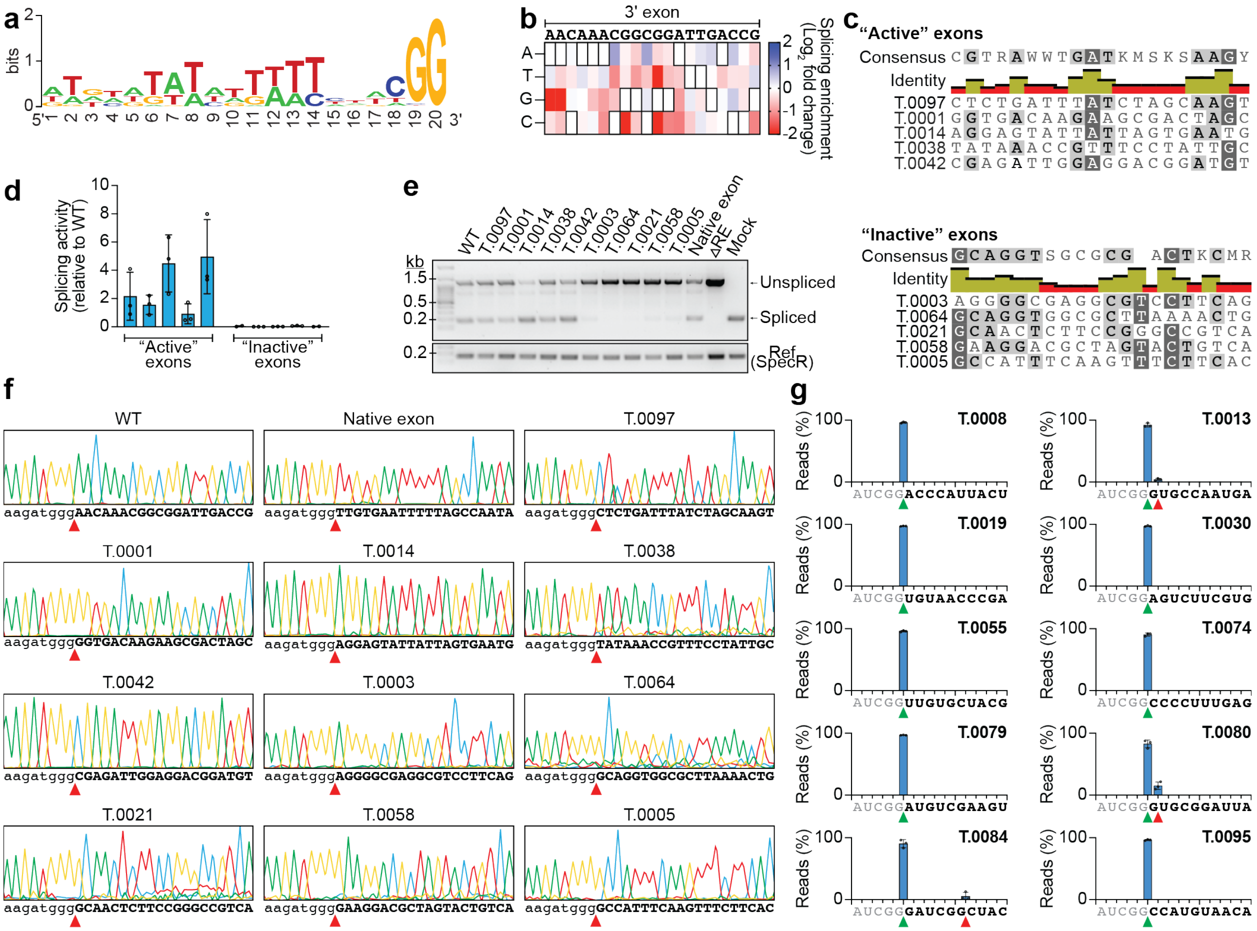


**Figure S6 | Effects of 3' exon sequence on splicing efficiency and splice site selection.**

**a)** Sequence logo revealing consensus motif of IStron RE sequences directly upstream of the alternative 3'SS selected. Logo was generated using WebLogo, with each variant 3' RE portion represented once.

**b)** Heatmaps depicting $log_2$-fold change (relative to wild-type) in the splicing assay for each nucleotide substitution across the 3' exon sequence. WT nucleotide identities contain black borders.

**c)** Sequence alignment of the 5 most active and the 5 most inactive exon sequences from the splicing assay. Alignments were done using MAFFT alignment; a consensus sequence is shown on top.

**d)** RT-qPCR analysis of splicing efficiency of IStrons with different 3' exon sequences. 'Active' and 'inactive' designations correspond to their abundance in library-scale splicing assays. All efficiencies are normalized to the WT 3' exon used in the library.

**e)** Agarose gel electrophoresis of RT-PCR products from splicing assays shown in **a**, showing unspliced and spliced products (top) relative to reference amplicons for a SpecR drug marker (bottom). -C indicates an IStron variant containing a scrambled RE sequence. Also included is a variant with the WT IStron RE sequence and its native 3' exon sequence from the native locus in *C. botulinum* strain BKT015925.

**f)** Sanger sequencing for all variants tested in this experiment (corresponding to data shown in Fig. 4D). The sequence of the spliced transcript is written below; the 3' exon sequence is in bold; red triangles denote splice sites.

**g)** Distribution of reads from library-scaled splicing experiments of variants with alternative 3' exon sequences. For each library member, the transcript backbone was analyzed to determine the exact splice site. Splice sites were classified as either WT splice site if the IStron library member spliced accurately at the end of the member, or alternative splice site if not. The most abundant alternative splice site was identified for each member. Shown are randomly selected variants from this pool. Library variant accessions are at the top right of the graph. Nucleotide sequences are shown below each distribution. Grey lettering denotes IStron RE sequence; bold lettering denotes the 3' exon; green triangle denotes WT splice site; red triangle denotes alternative splice site.
